# Cooperativity in Plant Plasma Membrane Intrinsic Proteins (PIPs): Mechanism of Increased Water Transport in Maize PIP1 Channels in Hetero-tetramers

**DOI:** 10.1101/239780

**Authors:** Manu Vajpai, Mishtu Mukherjee, Ramasubbu Sankararamakrishnan

## Abstract

Plant aquaporins (AQPs) play vital roles in several physiological processes. Plasma membrane intrinsic proteins (PIPs) belong to the subfamily of plant AQPs and they are further divided into two closely related subgroups PIP1s and PIP2s. Members of the two subgroups have been shown to have different transport properties. While PIP2 members are efficient water channels, PIP1s from some plant species have been shown to be functionally inactive. Aquaporins form tetramers under physiological conditions. PIP2s can enhance the water transport of PIP1s when they form hetero-tetramers. However, the role of monomer-monomer interface and the significance of specific residues in enhancing the water permeation of PIP1s have not been investigated at atomic level. We have performed all-atom molecular dynamics (MD) simulations of homo-tetramers of ZmPIP1;2 and ZmPIP2;5 from *Zea mays* and hetero-tetramers containing ZmPIP1;2 and ZmPIP2;5 with different stoichiometries and configurations. ZmPIP1;2 in a tetramer assembly will have two interfaces, one formed by transmembrane segments TM4 and TM5 and the other formed by TM1 and TM2. We have analyzed channel radius profiles, water transport and potential of mean force profiles of ZmPIP1;2 monomers with different types of interfaces. Results of MD simulations clearly revealed that TM4-TM5 interface and not the TM1-TM2 interface is important in modulating the water transport of ZmPIP1;2. We generated *in silico* mutants of specific residues that are involved in contacts with adjacent monomers. MD simulations of mutant tetramers highlighted the importance of I93 residue from the TM2 segment of ZmPIP2;5 for the increased water transport in ZmPIP1;2.

## Introduction

Plant aquaporins constitute an important component in the superfamily of Major Intrinsic Proteins (MIPs) ^1–2^. They are diverse and large number of plant MIPs have been identified in several plants including *Arabidopsis thaliana* ^3^, *Zea mays* ^4^, *Populus tricocarpa* ^5^, *Oryza sativa* ^6^, *Gossypium hirsutum^7^, Solanum lycopersicum* ^8^ and many others. Plant MIPs have been shown to be involved in several important physiological processes. They play a major role in regulating the plant-water homeostasis ^9^ and thus are instrumental in responding and adapting to various types of abiotic stress ^10–11^. Aquaporins in plant root and shoot mediate tissue hydraulics and can impact transpiration in plants ^12–14^ Key roles of plant aquaporins in other important plant-related function include nutrient transport, seed germination and emergence of lateral roots ^12, 15–16^. Apart from water, plant MIPs have been shown to transport diverse neutral solutes such as glycerol, carbon dioxide, hydrogen peroxide, urea, ammonia, silicic acid, arsenite and boron and ^1, 17–18^. With large number of plant MIPs identified, phylogenetic analyses have contributed significantly to identify and understand the different subgroups within plants and characterize them ^19^. Plant MIPs can be largely classified into five major groups: plasma membrane intrinisic proteins (PIPs), tonoplast intrinsic proteins (TIPs), nodulin 26-like intrinsic proteins (NIPs), small and basic intrinsic proteins (SIPs) and X-intrinsic proteins (XIPs) ^3–5^. They differ in subcellular localization, transport properties and stress response^2, 20^. Subcellular localization of several plant MIPs has been determined to be the plasma membrane (PIPs, XIPs and NIPs) ^21–23^, tonoplast (TIPs)^24^, nodule (NIPs)^25^ and endoplasmic reticulum (SIPs)^26^. In terms of transport, NIPs have been shown to transport silicon and boron ^27–28^. A few PIPs seem to be involved in carbon dioxide permeability ^29–30^. TIP members facilitate urea and ammonia transport across the membrane ^31–32^.

Three-dimensional structures of at least two plant MIPs have been determined. X-ray structures of spinach plasma membrane aquaporin SoPIP2;1 ^33^ and tonoplast intrinsic protein from *Arabidopsis thaliana* AtTIP2; 1 ^34^ show that the plant aquaporins adopt a similar hourglass helical bundle fold found in other MIP superfamily members across the three kingdoms ^35^. Under physiological conditions MIP members form homotetramers with each monomer forming a functional channel.

Among the plant MIP subfamilies, PIP subfamily members are one of the most extensively studied group. They are highly conserved and are further classified into two groups phylogenetically, namely, PIP1s and PIP2s. Sequences of plant PIPs are highly conserved. For example, the sequence identity between PIP members of maize varies between 64 to 100% ^4^ However, members belonging to PIP1 and PIP2 subgroup differ in many aspects including subcellular localization and transport properties. Among the two groups, mainly members belonging to PIP2 subgroup have been shown to be involved in water transport ^36–37^ In the case of PIP1s, reports regarding their ability to transport water are not conclusive. While in some species they are shown to be efficient water channels ^13, 38–40^, other PIP1 members exhibit relatively low water transport ^41–42^. In plants like maize, PIP1s do not seem to increase the osmotic water permeability coefficient (*P_f_*) prompting the authors to conclude that these channels are non-functional ^43–44^.

Several experimental studies have demonstrated that when PIP1 members are co-expressed with PIP2s, an increase in *P_f_* is observed. This was first shown in maize PIP1 and PIP2 members and Fetter et al. concluded that PIP1 and PIP2 members physically interact with each other resulting in hetero-tetramerization of both isoforms ^45^. Using confocal and FRET/fluorescent lifetime imaging microscopy, it has been shown that maize PIP2s help in relocalizing the PIP1s from endoplasmic reticulum to plasma membrane ^21^. Interactions involving the C-terminal part of loop E and possible disulfide bond between monomers formed by a cystine residue from loop A are some of the regions/residues that could potentially impact the formation and stability of such hetero-oligomers ^45–47^ The role of PIP1s as a modulator of membrane water permeability has also been demonstrated in PIPs of garden strawberry ^48^.

To find out whether there is any preference in the stoichiometry of PIP1-PIP2 hetero-tetramers, BvPIP1;1 and BvPIP2;2 from *Beta vulgaris* have been co-expressed in *Xenopus* oocytes. It has been shown that all hetero-tetrameric configurations (3:1, 2:2, 1:3) exist and all produced active channels localize in plasma membranes with equivalent efficiency in transporting water ^49^. Similarly, a large number of mutants have been generated for the *Arabidopsis* PIP2;1 channel and certain residues within and between monomers have been found to be important for the heteromer formation and for the trafficking of the protein from endoplasmic reticulum to plasma membrane ^50^. Modeling studies involving ZmPIP1;2 and ZmPIP2;5 hetero-tetramers have identified residues occurring at the monomer-monomer interface ^51^. Mutation of many of these interface residues either inactivates or activates the individual channels within the oligomeric assemblies. The role of specific residues at the monomer-monomer interface in the water transport and subcellular localization has been established in these mutagenesis studies.

Experimental studies have shown the importance of hetero-tetramer formation for the trafficking of PIP1 and PIP2 isoforms from ER to plasma membrane. Individual residues at the interface seem to have significant contribution in the heteromer formation and in the increased water transport of PIP1s. However, this phenomenon has not been investigated at atomic level. In this paper, we have performed extensive molecular dynamics simulations of PIP1 and PIP2 tetramers in explicit lipid bilayers. We have investigated two types of tetramers: homotetramers of PIP1s and PIP2s and hetero-tetramers involving both the PIP1s and PIP2s with different stoichiometries. For each type of tetramer, we have analyzed the channel properties, water transport and inter-monomer interactions. Based on our results, we have also generated *in silico* mutants of residues at the interface and compared with the wild-type tetramers. Our studies demonstrate that the interface formed by TM4-TM5 region of PIP1 can modulate the water transport when it interacts with PIP2. However, the TM1-TM2 interface of PIP1 does not seem to influence the function of the channel even if it interacts with PIP2 in hetero-tetramers.

## Materials and Methods

### Initial structures of homo- and hetero-tetramers of ZmPIP1;2 and ZmPIP2;5 channels

Since Chaumont and his colleagues used maize PIPs, ZmPIP1;2 and ZmPIP2;5, in their mutation studies to demonstrate that PIP1 transport is modulated by PIP2 ^51^, we considered the sequences of the same maize PIPs. The UniProt ^52^ accession IDs corresponding to ZmPIP1;2 and ZmPIP2;5 are Q9XF59 and Q9XF58 respectively. Experimental structures of 12 unique MIP channel structures have been determined from diverse species. MIP structures from different sources such as yeast, *Escherichia coli*, *Plasmodium falsiparum*, archaea, plants and mammals, exhibit a conserved hourglass helical fold^35^. Using homology modeling approach, we have constructed three-dimensional models of the monomers of more than 1400 MIP channels and these structural models are available from the MIPModDB database ^53^. MIP homology models from diverse species were built using three template structures determined experimentally from *E. coli* (PDB ID: 1FX8), archaea (PDB ID: 2F2B) and bovine (PDB ID: 1J4N). Details of homology modeling protocols are described in detail in our earlier publications ^5, 35, 53–56^. Homology models of ZmPIP1;2 and ZmPIP2;5 monomers were downloaded from MIPModDB ^53^. We also calculated the root mean square deviation (RMSD) of the models with the plant spinach aquaporin X-ray structures SoPIP2;1 (PDB IDs: 1Z98 and 2B5F) and they are found to be very close to the experimentally determined structures. RMSD between ZmPIP1;2 and ZmPIP2;5 models and the SoPIP2;1 crystal structures are between 0.7 to 0.8 Å giving enough confidence to proceed with the modeled structures. MIP channels are found to exist as tetramers under physiological conditions and hence the same set of transformations that are used to generate the biological assembly of experimentally determined structures were applied on ZmPIP1;2 and ZmPIP2;5 monomer models to generate tetramers. This procedure was followed to obtain ZmPIP1;2 and ZmPIP2;5 homo- and hetero-tetramers with varying stoichiometries and configurations.

A total of six unique configurations were considered (Table 1). Two of them were homo-tetramers where all monomers were either ZmPIP1;2 or ZmPIP2;5. The remaining four consisted of different combinations of ZmPIP1;2 and ZmPIP2;5. Each tetramer thus generated was first energy minimized using GROMACS 4.5.6 ^57^. We have also investigated mutants in which point mutations were introduced at the monomer-monomer interface. Dunbrack rotamer libraries ^58^ and UCSF Chimera ^59^ were used for this purpose.

**Table 1:** Different stoichiometries and configurations of ZmPIP1;2 and ZmPIP2;5 tetramers and their water transporting properties

| ZmPIP tetramer<sup>a</sup> | PIP1:PIP2<sup>b</sup> | Water permeation events<sup>c</sup> | Osmotic permeability ($p_f$)<sup>d</sup> |
| --- | --- | --- | --- |
| ●●<br>●● | 4:0 | 14.0 | 1.16 |
| ●●<br>○● | 3:1 | 98.1 | 3.02 |
| ●○<br>○● | 2:2 (d) | 68.2 | 2.46 |
| ●●<br>○○ | 2:2 | 58.6 | 2.25 |
| ●○<br>○○ | 1:3 | 98.2 | 3.12 |
| ○○<br>○○ | 0:4 | 64.0 | 2.26 |

<sup>a</sup>ZmPIP1;2 and ZmPIP2;5 are represented as “●” and “○” respectively. Different combinations of ZmPIP1;2 and ZmPIP2;5 varying in stoichiometries and configurations are shown.
<sup>b</sup>The ratio of ZmPIP1;2 and ZmPIP2;5 in a given tetramer is shown. The ratios 4:0 and 0:4 represent homotetramers of ZmPIP1;2 and ZmPIP2;5 respectively. Both the heterotetramers 2:2(d) and 2:2 have the same number of ZmPIP1;2 and ZmPIP2;5 monomers but differ in their configurations. The two ZmPIP1;2 (or ZmPIP2;5) monomers are present diagonal to each other in 2:2(d) while in 2:2 they are adjacent.
<sup>c</sup>Average number of water permeation events calculated for all the monomers of a given tetramer from two independent simulations is given.
<sup>d</sup>Average osmotic permeability calculated for all the monomers of a given tetramer from two independent simulations is given. Osmotic permeability is calculated as described in the Methods section. Osmotic permeability is in the units of $10^{-14}$ cm<sup>3</sup>/s.

### Molecular dynamics simulations of ZmPIP1;2 and ZmPIP2;5 tetramers

Each of the six tetramers thus generated was first solvated in a pre-equilibrated POPE bilayer patch (400 POPE lipids) ^60^ and we followed the protocol proposed by Kandt et al. ^61^. Using this approach, we first expanded the pre-equilibrated POPE bilayer within XY plane and those lipids within a specific cut-off distance from the tetramer were deleted. The initial dimension was obtained by compressing the box size using a series of scaling steps followed by energy minimization. Berger lipid parameters ^62^ and OPLS-AA force field for proteins ^63^ were used for minimization and subsequent simulations. The lipid-embedded systems were further solvated with TIP3P ^64^ waters and neutralized with appropriate number of counter-ions. Each system was initially simulated for a period of 100 ps with NVT (constant number of atoms, volume and temperature) ensemble and harmonic restraints of 10000 kJ/mol/nm^2^ were imposed on protein atoms and headgroups of lipids. We carried out another 700 ps equilibration in seven stages and the lipid headgroup restraints were gradually removed during this time. Further equilibration of 1 ns was performed in NPT (constant number of atoms, pressure and temperature) ensemble by retaining the restraints on the protein atoms. Parinello-Rahman barostat ^65^ was used to maintain a constant pressure of 1 bar. The systems were simulated at a temperature of 310 K and the temperature was maintained using Nose-Hoover coupling ^66^. Finally, another 15 ns equilibration was carried out without any restraints. This was followed by the production run of 200 ns and all the analyses were performed on the trajectories generated during these production runs. Another independent simulation for each system for a period of 200 ns production run was performed by changing the initial velocities and following the same protocol. Thus a total of 2.4 μs (6 systems each simulated for a period of 200 ns and two independent simulations for each system) production runs were performed on six different tetramers of maize PIP aquaporins. Additionally, another 0.8 μs production runs were performed on the mutant systems of maize PIPs.

### Potential of mean force (PMF) profiles of water permeation

PMF profiles of the permeation of water molecules for each system were calculated using the approach suggested by de Groot and Grubmuller ^67^. Number of channel water molecules residing within the PIP monomers was found out from each MD simulated structure saved at every 10 ps using our in-house scripts. Probability distribution of water molecules was calculated for the entire 200 ns trajectory for each simulation. This information was used to calculate the PMF [G_i_(z)] profile using the following equation:

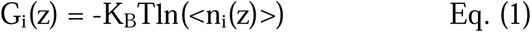

Where K_B_ and T are respectively the Boltzmann constant and temperature and <n_i_(z)> is the average number of water molecules as a function of channel axis. We also calculated PMF profiles of more than one monomer of the same type (see below). In these cases, the MD trajectories of individual monomers were merged and the average PMF profile was calculated as described above.

### Water permeation

The number of water permeation events and osmotic permeability (*p_f_*) were calculated to quantify the water transport of the PIP monomers. The positions of water molecules within the channel were ascertained every 1 ps. We generated bins for every 200 ps and calculated the mean-squared displacement for the water file which was then used to calculate the osmotic permeability (*p_f_*) as described by Zhu et al. ^68^. The *p_f_* value was obtained from the diffusion constant using the following equation.

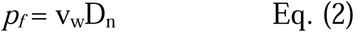

Where v_w_ is the volume of a single water molecule and D_n_ is the diffusion constant calculated using Einstein equation from the mean-squared displacement of the water file.

For each monomer, number of water permeation events was calculated by noting the number of water molecules which completely permeated the channel during the course of the simulation using our in-house scripts.

### Temporal channel radius profiles

For all the simulated systems, channel radius profiles for all the monomers were calculated using the program HOLE ^69^. HOLE typically calculates channel radius profile for a given channel protein and the radius profile can be plotted as a 1-dimensional diagram. In the present analysis, we calculated channel radius profiles by considering each monomer from the MD snapshot extracted every 10 ps. We have generated two-dimensional channel radius profiles in which the evolution of channel radius as the function of time is plotted at each point along the channel axis. The channel radius profiles were averaged over 2 ns bins at each point while generating the 2D-profile. We refer this as “temporal channel radius profile” reflecting the fact that the 2D-plot has information about time evolution of channel radius at points along the channel axis. We have also calculated the temporal channel radius profiles for a set of monomers having similar interfaces. In these cases, the averaging at each bin was done over all the trajectories of individual monomers. The 2D-profile thus generated was then plotted for the entire length of the simulations.

### Contact maps

Distances between heavy atoms of interacting monomers that are part of the tetramers formed either by wild-type PIPs or in systems in which one of the PIP monomers was mutated *in silico* were calculated using the MDTraj Python module ^70^. If the distance between two heavy atoms of interacting monomers is within 4.0 Å during at least 70% of the simulation time, then these two atoms are assumed to have contacts. Contact maps were plotted by summing and normalizing the contacts between the transmembrane segments of interacting monomers. We have also generated contact difference maps to ascertain differences between the contacts formed by monomers which differ in the interfacial region.

### Essential dynamics analysis

Essential dynamics analysis, also known as Principal Component Analysis (PCA), was performed on individual monomers using the built-in analysis tool of GROMACS. When we considered the monomers with the same type of interface, we concatenated their trajectories to perform PCA on the complete set. In each case, we observed that more than 90% of the motions were accounted by the first 10 principal components. The directions and the extent of fluctuations in the transmembrane segments of PIP1 were visualized using the extremea of the first principal components with the help of PyMol viewer^71^.

## Results

The mechanism of water permeation and glycerol selectivity has been investigated in aquaporin and aquaglyceroporin using molecular dynamics approach ^67, 72–73^. To investigate the differences in the water transporting properties of homo- and hetero-tetramers comprising PIP1 and PIP2 monomers, we have first performed molecular dynamics simulations of six oligomers with different combinations of ZmPIP1;2 and ZmPIP2;5. For each system, two independent simulations each for a period of 200 ns were carried out. Water permeation events, osmotic permeability, temporal channel radius profiles and potential of mean force are some of the properties analyzed to find the channel behavior of individual monomers. Analyses of interfacial contacts and essential dynamics motions were performed to understand the influence of neighboring monomers on the water transport. Table 1 summarizes the tetramers considered and the transport properties of both homo- and hetero-tetramers. Water permeation events and osmotic permeability of an individual monomer from two independent simulations for all the simulated systems are provided in the Supplementary Information (Figure S1 and Figure S2). As reported in the experimental studies ^49^, the homotetramers formed by ZmPIP1;2 resulted in the lowest water transport as observed by the average number of water permeation events and the osmotic permeability calculated by combining the two independent simulations and all monomers. When the same data was compared for the homotetramers of ZmPIP2;5, there is a significant increase in the water permeability, again agreeing with the previously reported experimental studies ^49^. All four hetero-tetramers exhibit higher water permeability compared to the ZmPIP1;2 homo-tetramers. Water-transporting efficiency of two of the hetero-tetramers has surpassed even the ZmPIP2;5 homo-tetramers indicating that the ZmPIP1;2 in these oligomers are involved in higher water transport than that found in the homo-tetramers of ZmPIP1;2. The osmotic permeabilities of two of the ZmPIP2;5 heterotetramers (3:1 and 1:3) are comparable to the recently reported *p_f_* value of the water transporting channel AQP5 obtained from MD simulations ^74^.

This brings into the question the capability of ZmPIP1;2 in transporting more water molecules when it is present as part of a hetero-tetramer in comparison to homo-oligomers. In the tetramer assembly, each monomer interacts with two adjacent monomers and hence has two interfaces. The transmembrane helices TM4 and TM5 form one interface while TM1 and TM2 form another interface through which interactions with the neighboring monomers take place (Figure 1). In homotetramers, all the monomers are of the same type and hence the interfaces formed by them do not differ from one monomer to another one. However in a hetero-tetramer, at least one monomer will have interfaces formed by two different PIP subtypes. We have investigated the transport properties of ZmPIP1;2 in both kinds of environments by considering the effects of neighboring interactions of monomers in homo- and hetero-tetramers.

**Figure 1:**
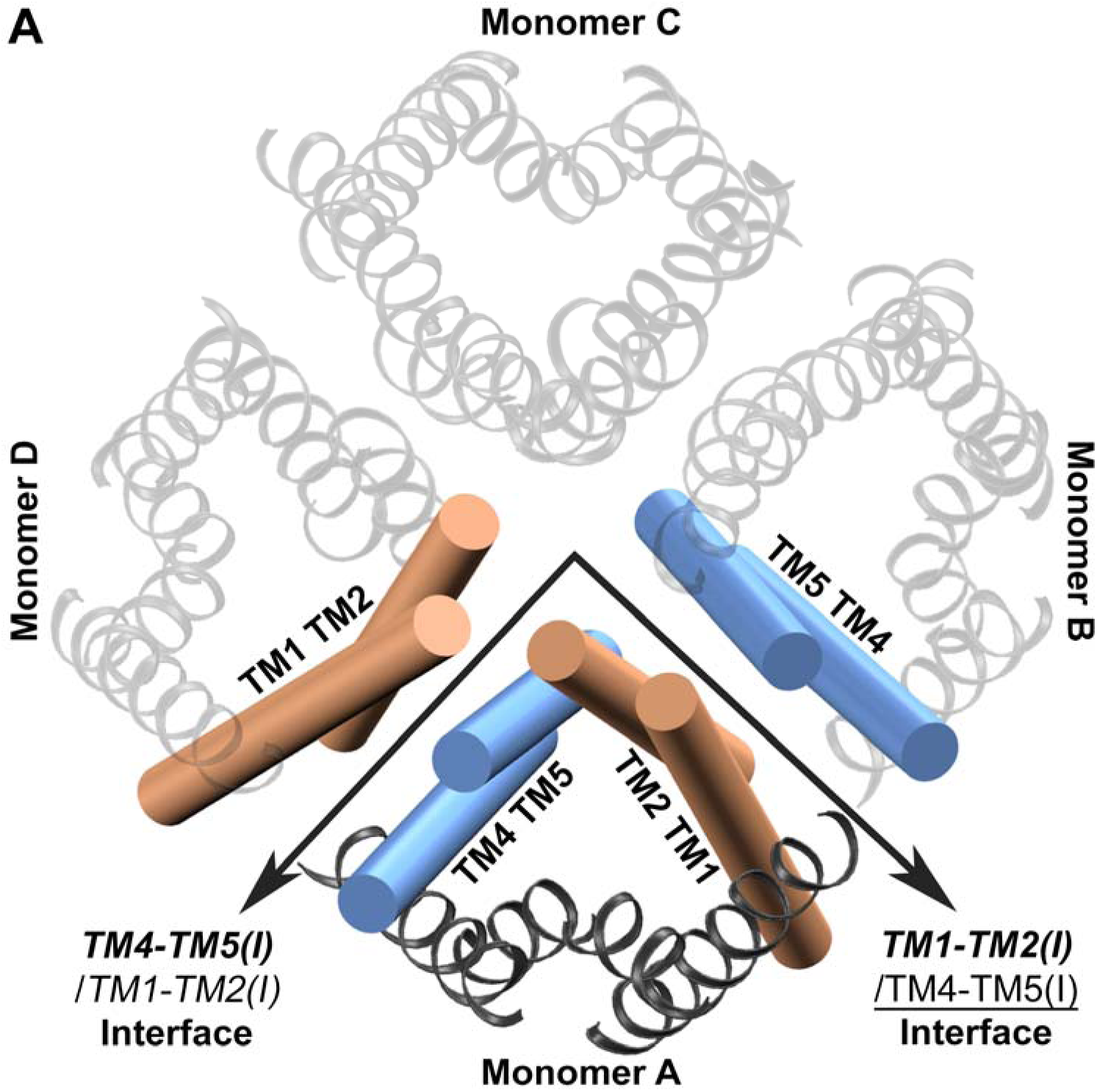
ZmPIP1;2 homo-tetramer showing two different monomer-monomer interfaces formed by TM1-TM2 (orange) and TM4-TM5 (blue) segments. The monomer under consideration is shown in opaque and the other three monomers are displayed as transparent. The convention for naming the interfaces is explained in the main text. Unless otherwise stated, the molecular plots in this figure and subsequent figures were generated using VMD software ^75^.

### Interfaces of ZmPIP1;2 in homo- and hetero-tetramers

There are four possible interface pairs for any given ZmPIP1;2 monomer in a homo- or hetero-tetrameric configuration and they are as follows.

Type I: Both the interacting neighbors are ZmPIP1;2. In this case, the interfaces will be ***TM4-TM5(I)****/TM1-TM2(I)* and ***TM1-TM2(I)****/*<u>TM4-TM5(I).</u>

Transmembrane segments of monomer under consideration are depicted in bold and those of adjacent interacting neighbors are either shown in italics (when in the interface is formed with TM4-TM5 helices) or underlined (when the TM helices interact with TM1-TM2 segments). The Roman (I) or (II) indicates ZmPIP1;2 or ZmPIP2;5 respectively. This notation will be followed throughout the paper.

Type II: The TM4-TM5 helices of ZmPIP1;2 interacts with TM1-TM2 of ZmPIP2;5 while TM1-TM2 helices of ZmPIP1;2 interacts with the neighboring TM4-TM5 segments which is also ZmPIP1;2. The notation for the interface in this case is:

***TM4-TM5(I)****/TM1-TM2(II)* and ***TM1-TM2(I)****/*<u>TM4-TM5(I)</u>

Type III: The TM4-TM5 helices of ZmPIP1;2 interact with TM1-TM2 of ZmPIP1;2 while TM1-TM2 helices of ZmPIP1;2 interact with the neighboring TM4-TM5 of ZmPIP2;5. The interfaces in this case will be denoted as:

***TM4-TM5(I)****/TM1-TM2(I)* and ***TM1-TM2(I)/***<u>TM4-TM5(II)</u>

Type IV: Both the interacting neighbors are ZmPIP2;5. The interfaces here are represented as:

***TM4-TM5(I)****/TM1-TM2(II)* and ***TM1-TM2(I)****/*<u>TM4-TM5(II)</u>

For the populations of monomers with different types of interfaces described above, we have compared water permeation events, osmotic permeability, PMF profiles and channel radius profiles to find out the influence of interacting neighbors on water transport. Analyses of the monomers with different types of interfaces are discussed below.

### ZmPIP1;2 monomers: Type-I category

All the monomers within the homo-tetramers of ZmPIP1;2 belong to Type-I category and they show minimum water transport (on an average only 14 water permeation events) amongst all the systems studied. The only other monomer with this type of interface is in the tetramer with 3:1 combination of ZmPIP1;2 and ZmPIP2;5 (Table 2). In this system, the adjacent interacting monomers are ZmPIP1;2 while the lone ZmPIP2;5 is located diagonal to this monomer. Overall, the 3:1 hetero-tetramer is one of the highest water-transporting systems with average number of water permeation nearly 98 and osmotic permeability 3.14. However, the monomer with Type-I interface in this system shows reduced permeation events and *p_f_* value indicating that the interacting interfaces from the neighboring monomers make it behave almost like a monomer in ZmPIP1;2 homo-oligomer. It provides the first evidence that the interfaces with the neighboring monomers influence the water transport. To further elucidate the factors responsible for the reduced water transport, we have plotted the temporal channel radius profiles calculated for the monomer under investigation. This property was compared with that of ZmPIP1;2 homo-tetramers which shows that the channel is severely constricted near the aromatic/arginine selectivity filter throughout the simulation time in the homo-oligomer (Figure 2A). The same radius profile analysis of the monomer in the 3:1 system with the Type-I interface displays a profile similar to that of ZmPIP1;2 homotetramer showing that both ar/R selectivity filter and the region near the NPA motif are very narrow preventing the water molecules to pass these regions (Figure 2B).

**Table 2:**
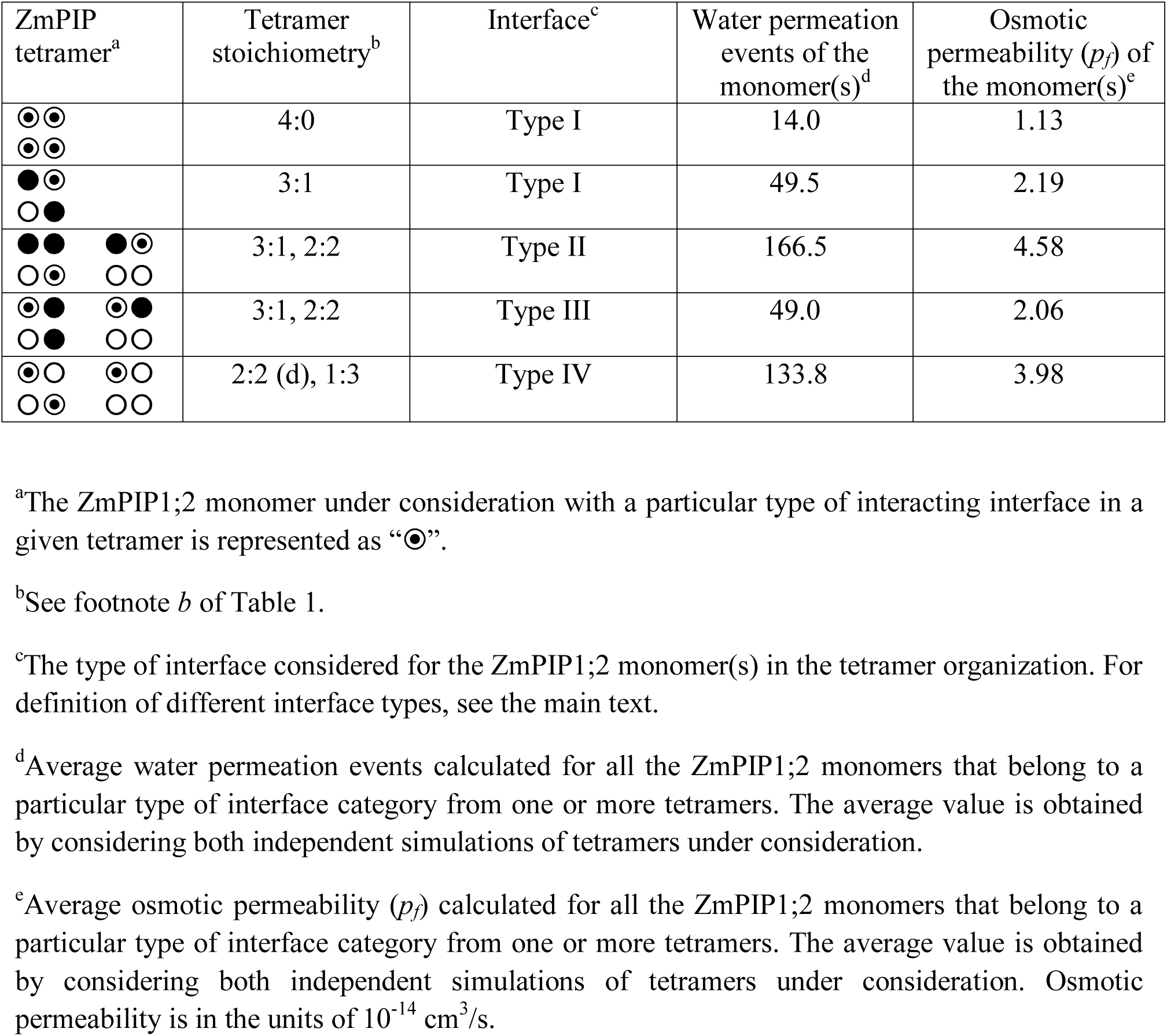
Water transport properties of ZmPIP1;2 monomers with different interacting interfaces with adjacent monomers

**Figure 2:**
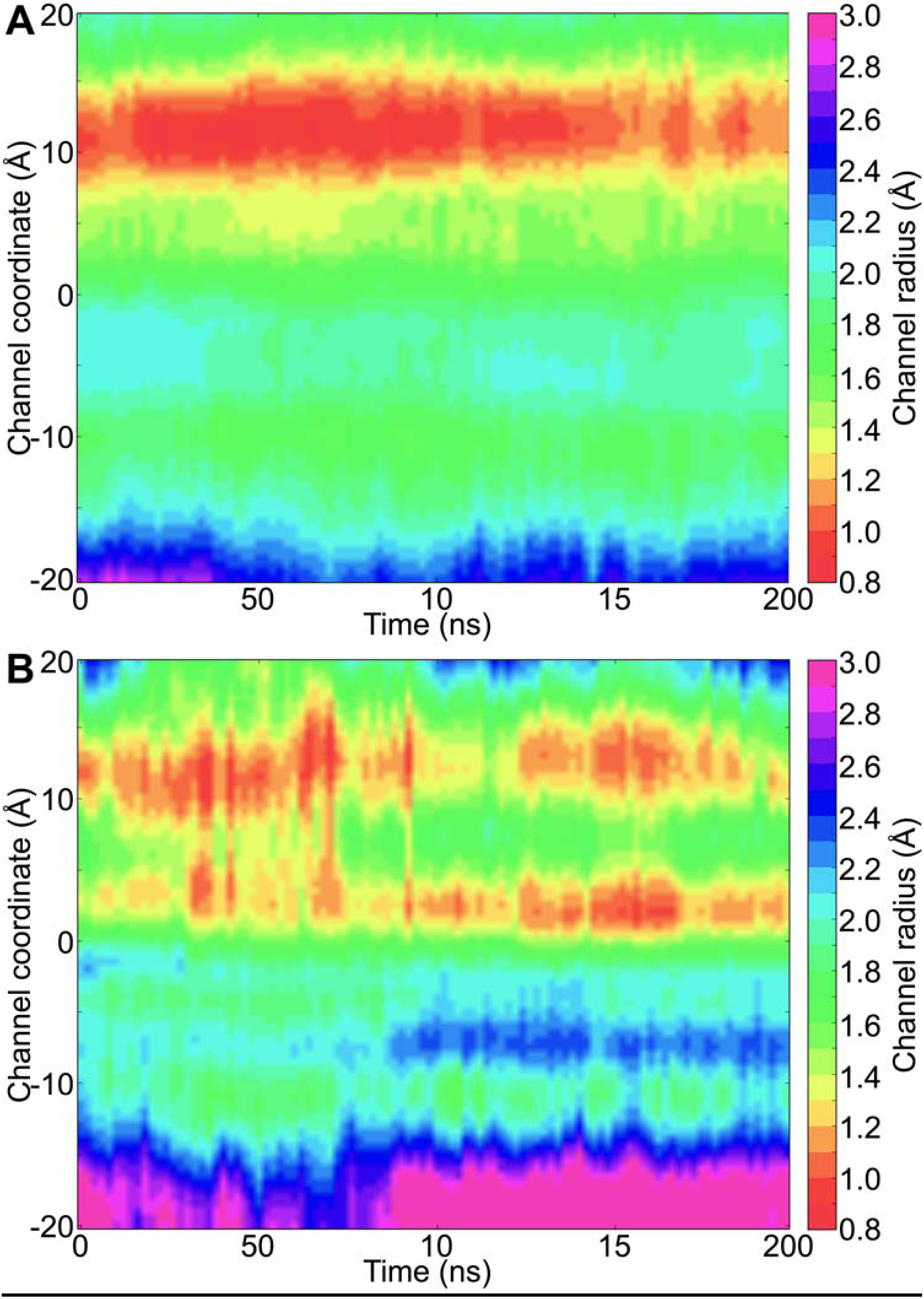
Temporal channel radius profiles of (A) ZmPIP1;2 homo-tetramer and (B) ZmPIP1;2 monomer from Type-I category in the 3:1 hetero-tetramer system. The point ‘0’ along the channel axis represents the NPA region. The ar/R selectivity filter is located at approximately 10 Å. Each point along the channel axis is the average calculated from 2 ns bin for all the monomers considered. For example, for the homo-tetramer system, all four monomers from the two independent simulations (total 8 monomers) were considered and a point along the channel axis in this 2D-profile is an average over 8 monomers in a 2 ns bin at that time.

### ZmPIP1;2 monomers: Type II category

There are two types of tetramers (3:1 and 2:2) in which a ZmPIP1;2 has interfaces that belong to Type II category. In this group of monomers, TM4-TM5 of ZmPIP1;2 interacts with TM1-TM2 of ZmPIP2;5 whereas the other interface is formed with another ZmPIP1;2 and there is one such monomer in both hetero-tetramers. The average water permeation events of the monomers of Type II group in 3:1 and 2:2 systems is the highest among all the monomers indicating that the PIP1:PIP2 interface of this type has a greater role in efficient water transport. The same is also reflected in the osmotic permeability (Table 2). This also explains as to why the overall water transport of both tetramers, 3:1 and 2:2, is higher than that of homotetramer 4:0. To further illustrate the increase in the water transport, we have plotted the temporal channel radius profiles calculated for both the monomers with the Type II interface from 3:1 and 2:2 tetramers (Figure 3A). Although initially the channel was narrow in the ar/R selectivity filter region till 50 ns, it becomes wider for the rest of the simulation. This can explain why the monomers from the Type II group could conduct large number of water molecules across the channel.

**Figure 3:**
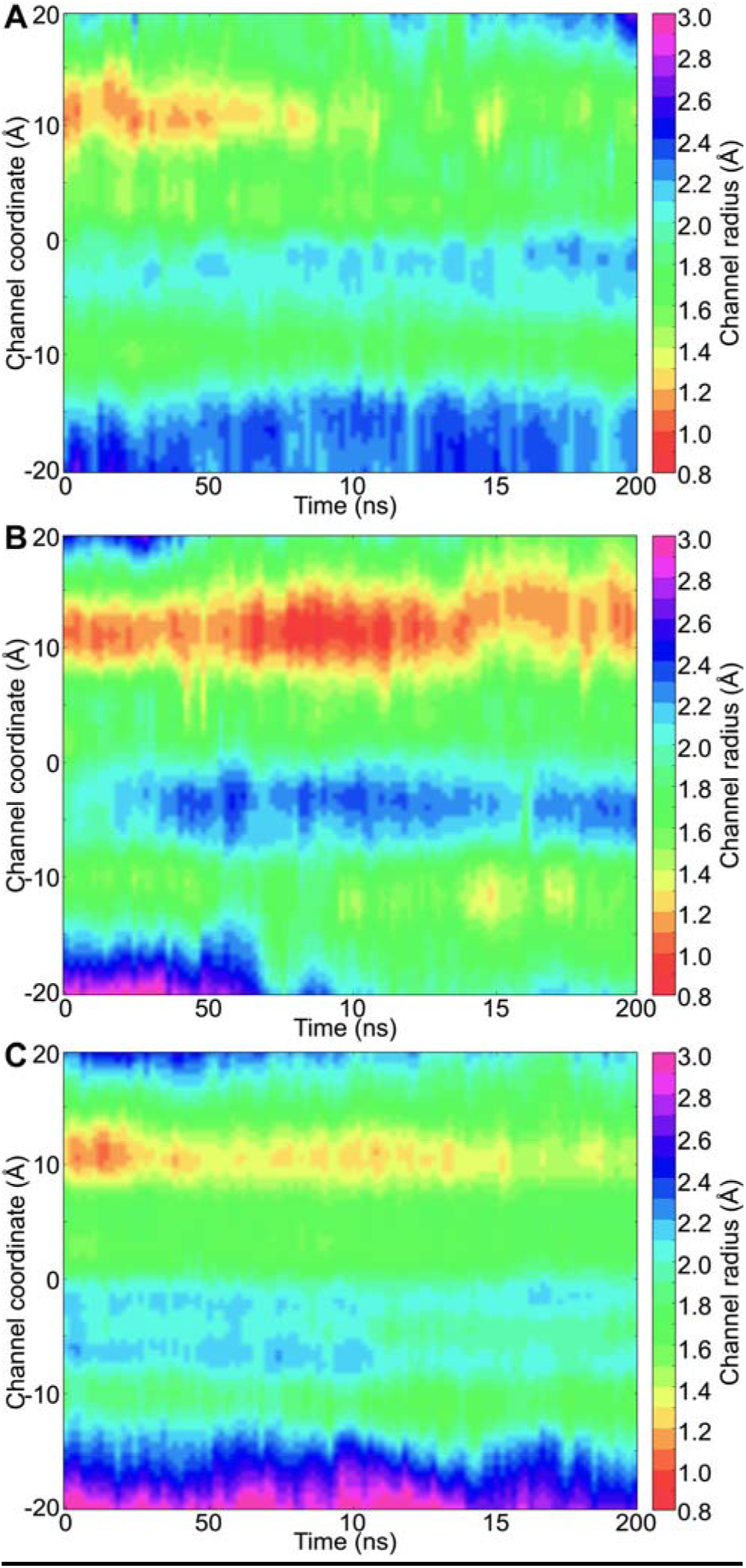
Temporal channel radius profiles of ZmPIP1;2 monomers (A) Type II, (B) Type III and (C) Type IV categories from hetero-tetramers. The number of monomers used to generate the 2D channel radius profiles for each category is 4, 4 and 6 respectively for Type II, Type III and Type IV. For all other details, see caption of Figure 2 and Table 2.

### ZmPIP1:2 monomers: Type III category

In this type, the TM1-TM2 side of the ZmPIP1;2 monomer under consideration interacts with the TM4-TM5 of ZmPIP2;5 while the other interface is formed with another ZmPIP1;2 monomer. Two monomers with 3:1 and 2:2 tetramers have interfaces that belong to Type III group and the average water permeation events in these monomers is only 49 which is similar to the monomer in 3:1 system with Type I interface. This indicates that the interface formed by the interaction of TM1-TM2 of ZmPIP1;2 with the TM4-TM5 of ZmPIP2;5 does not influence the water transport of the ZmPIP1;2 monomer. In contrast, the interface formed between the TM4-TM5 of ZmPIP1;2 and TM1-TM2 of ZmPIP2;5 has the greatest influence in increasing the efficiency of ZmPIP1;2’s water transport as observed in the monomers from Type II group. This is also evident from the temporal channel radius profiles of ZmPIP1;2 monomers that belong to Type III group from 3:1 and 2:2 systems (Figure 3B). There will be clearly an energy barrier due to the narrow channel width (< 1.0 Å) near the ar/R selectivity filter region.

### ZmPIP1;2 monomers: Type IV category

Three monomers belong to Type IV group among all the configurations investigated and are present in 2:2(d) and 1:3 tetramers. In this category, both neighbors of ZmPIP1;2 are ZmPIP2;5 and thus both TM4-TM5 and TM1-TM2 regions of ZmPIP1;2 interact respectively with TM1-TM2 and TM4-TM5 of ZmPIP2;5. Average water permeation of the three monomers from the two systems is about 134 during the simulation period. This is the second highest rate of water transport next the ZmPIP1;2 monomers from Type II group. Corresponding *p_f_* value of 4.07 is also the second highest next only to the monomers belonging to Type-II category. This again can be attributed to the interaction of TM4-TM5 side of ZmPIP1;2 with the TM1-TM2 region of ZmPIP2;5. Analysis of temporal channel radius profiles also illustrates that the channel is relatively wider in the ar/R selectivity filter region compared to that found for the monomers from Type I or Type III categories (Figure 3C). This could explain why ZmPIP1;5 monomers from Type IV group conduct larger number of water molecules.

### TM4-TM5 interfacial region of ZmPIP1;2 monomers seems to be important for regulating water transport

The above analyses clearly demonstrate that whenever the TM4-TM5 of ZmPIP1;2 interacts with TM1-TM2 of ZmPIP2;5, there is a significant increase in the water permeability of ZmPIP1;2. Hence, we combined the data from all the monomers from Type II and Type IV categories. The average and standard deviation of the number of water molecules transported through the ZmPIP1;2 monomers of Type II or Type IV group is 147 ± 115. When we compare the monomers of Type I or Type III category in hetero-tetramers, they transport 49 ± 43 water molecules. When the same analysis was performed on ZmPIP1;2 homotetramers, they transport on an average of only 14 water molecules. Since the data is not normally distributed, we performed Mann-Whitney U test to find whether the difference observed in the water transport between the three categories of monomers has any significance. The difference between the homotetramers and the monomers from Type II/Type IV group is statistically extremely significant with *p* value 0.00062. However, the number of water permeation events between homotetramers and the monomers of Type I/Type III category from hetero-tetramers does not show any significance with *p* value 0.139. This implies that the permeation behavior of Type I/Type III monomers is similar to that of homotetramers further confirming that there is a preference for TM4-TM5 interface in modulating ZmPIP1;2 water transport. Temporal channel radius profiles consolidated for all the monomers from the two categories are shown in Figure 4A and Figure 4B. We can see that the Type I/Type III monomers and the monomers from homotetramers (Figure 2A) have significant constriction near the ar/R selectivity filter. This constriction has largely disappeared in monomers from Type II/Type IV category.

**Figure 4:**
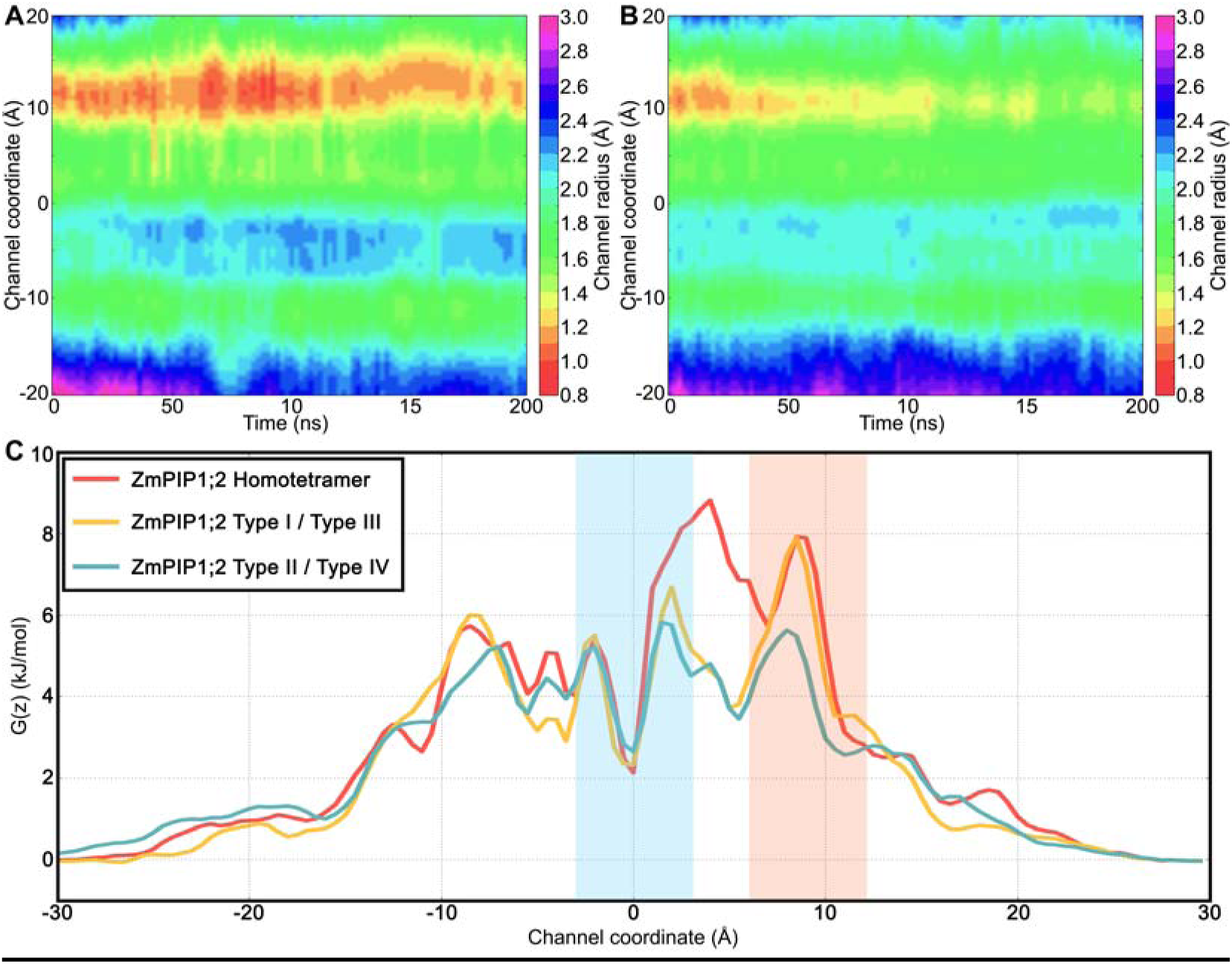
Temporal channel radius profiles of ZmPIP1;2 monomers consolidated for (A) Type I/Type III and (B) Type II/Type IV categories from different hetero-tetramer systems. A total of 6 and 10 monomers were considered for Type I/Type III and Type II/Type IV respectively. (C) Average PMF profiles calculated for ZmPIP1;2 monomers from homo-tetrameric system, ZmPIP1;2 monomers belonging to Type I/Type III and Type II/Type IV groups.

A similar exercise was carried out for ZmPIP2;5 monomers as well. We have considered the ZmPIP2;5 monomers whose TM4-TM5 region interacted with TM1-TM2 of ZmPIP1;2. Average water permeation events were found to be 31 ± 29. For all the ZmPIP2;5 monomers whose TM4-TM5 region interacted with the TM1-TM2 surface of another ZmPIP2;5, the average number of transported water molecules is 86 ± 58. When the ZmPIP2;5 monomers from the homotetramers were considered, the water permeability is 64 ± 59. We found that the difference in the water permeability between the three groups is not significant (*p* > 0.01). This implies that there is no significant change in water permeation of ZmPIP2;5 when it is present in a hetero-tetramer versus when it is present in homotetramer further implicating the regulatory role of TM4-TM5 interface only in the case of ZmPIP1;2.

To further demonstrate the importance of TM4-TM5 interface in the hetero-tetramers, we evaluated the potential of mean force (PMF) profiles of water transport for homo- and hetero-tetramers in which the TM4-TM5 interface of ZmPIP1;2s interacts with TM1-TM2 of either ZmPIP2;5 or ZmPIP1;2. As explained above, there are three categories of ZmPIP1;2 monomers in the context of ZmPrP1;2’s transport efficiency. ZmPIP1;2 monomers from Type II/Type IV group, ZmPIP1;2 monomers from Type I/Type III category and ZmPIP1;2 monomers within homo-oligomers. PMF profiles for all the three ZmPIP1;2 groups are plotted in Figure 4C. Average PMF profile calculated for the ZmPIP1;2 monomers from Type I/Type III group exhibit a larger energy barrier near the NPA motif and also in the ar/R selectivity filter region. ZmPIP1;2 monomers from the homo-tetramers also display a large energy barrier in the ar/R selectivity filter region. However, PMF profile of ZmPIP1;2 monomers from Type II/Type IV group indicates that the energy barrier in the same selectivity region has come down by 2 to 2.5 kJ/mol. Unlike Type I/Type III ZmPIP1;2 monomers, there is no peak next to NPA region for ZmPIP1;2 monomers from Type II/Type IV group. Both the number of instances of water permeation and PMF profiles show that the ZmPIP1;2 monomers from Type II/Type IV category are more efficient in transporting water molecules. This analysis reiterates that the interface involving interactions of TM4-TM5 from ZmPIP1;2 with TM1-TM2 of ZmPIP2;5 plays a major role in modulating the water transport properties of ZmPIP1;2.

For the purpose of comparison, we also plotted the PMF profiles of ZmPIP2;5 monomers belonging to all three categories (Figure S3 in Supporting Information). There is no major difference between the three types of ZmPIP2;5 monomers. This indicates that the interactions between ZmPIP1;2 and ZmPIP2;5 monomers in the hetero-tetramers at the TM4-TM5 interface of ZmPIP1;2 can enhance the water transport of ZmPIP1;2 only and similar phenomena may be absent in ZmPIP2;5. Although the TM1-TM2 interface of ZmPIP1;2 can also have interactions with ZmPIP2;5, our simulation studies have unambiguously demonstrated that TM4-TM5 interface is more important in modulating the water transport properties of ZmPIP1;2 monomers when they form hetero-tetramers with ZmPIP2;5.

### *In silico* mutants of residues at the TM4-TM5 interface

When we compared the sequences of ZmPIP1;2 and ZmPIP2;5, we found 64.4% sequence identity and 79.5% sequence similarity between these two sequences. The alignment in the TM1 and TM2 regions between these two sequences is shown in Figure 5A. Five positions in the transmembrane helical segments at the monomer-monomer interfacial region exhibit differences between ZmPIP1;2 and ZmPIP2;5 sequences. We generated *in silico* mutants of ZmPIP2;5 substituting these five residues corresponding to those from ZmPIP1;2 in the heteromeric system 2:2 (d). Two additional positions in the C-terminus of TM1 of ZmPIP2;5 were also mutated in this system to ascertain if the residues just outside the transmembrane region were also involved. The hetero-tetramer 2:2 (d) was chosen for the mutant system so that one ZmPIP2;5 will be mutated and the other will be the wild-type ZmPIP2;5 and can serve as a control. The interfacial regions at the TM4-TM5 of ZmPIP1;2 interacting with ZmPIP1;2 and ZmPIP2;5 are shown in Figure 5B and Figure 5C respectively. We first substituted these seven interfacial residues of ZmPIP2;5 corresponding to that of ZmPIP1;2 and the starting structure of this mutant system is shown in Figure 6A. In the wild-type 2:2 (d) tetramer, both ZmPIP1;2s will have interface that can be classified as Type IV. In the mutant 2:2(d), one ZmPIP1;2 will have interface similar to Type-III monomer. We performed two independent MD simulations of this mutated system following the same protocol used to simulate wild-type homo- and hetero-tetramers. We calculated the number of water permeability events and *p_f_* for all the monomers. The ZmPIP1;2 with Type IV category interface transported on an average 168 water molecules while the ZmPIP1;2 which acquired Type III-like interface in the mutant system transported only an average of 26 water molecules. In the wild-type 2:2(d) system, the average number of water molecules transported by both ZmPIP1;2 members is 87 and is comparable to Type IV ZmPIP1;2 monomer in the mutant system. The ZmPIP1;2 with Type III-like interface behaved more like ZmPIP1;2s in wild-type homotetramer or Type I/Type III ZmPIP1;2 in wild-type hetero-oiligomers. The corresponding *p_f_* values for ZmPIP1;2 with Type IV and Type III-like interfaces are 3.90 and 1.98 respectively confirming that the transport property of ZmPIP1;2 in Type III-like interface is compromised.

**Figure 5:**
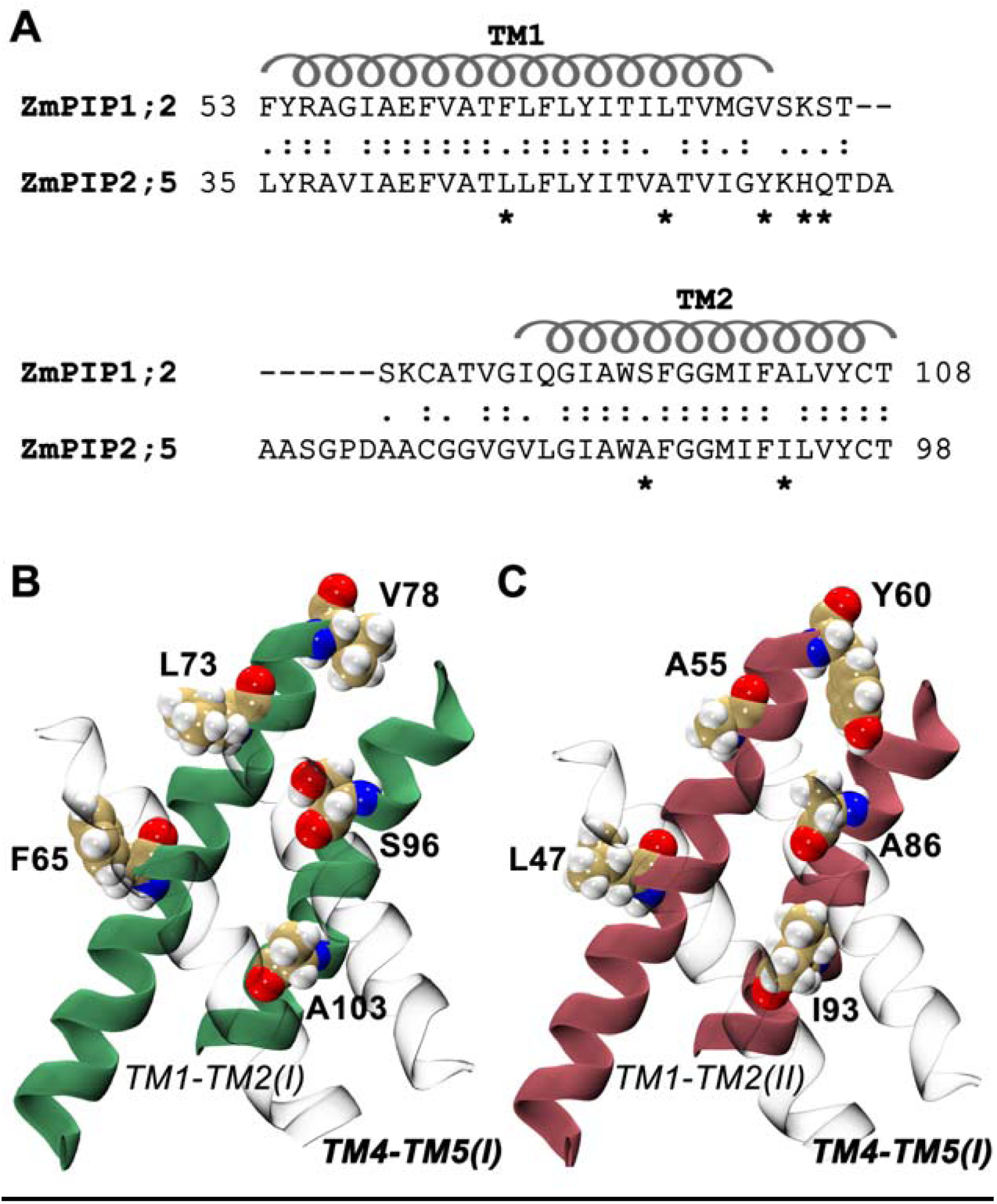
(A) Pairwise sequence alignment of TM1 and TM2 helical segments along with the linker region are shown for ZmPIP1;2 and ZmPIP2;5. Residues that are at the interface and are different between the two sequences are marked. The transmembrane helical regions are indicated. The seven marked positions are mutated in the 2:2(d) heteromer system and the mutant oligomer was simulated to understand the influence of the interfacial residues. Molecular plots showing the interfaces formed by TM4-TM5 of ZmPIP1;2 with TM1-TM2 of (B) ZmPIP1;2 and (C) ZmPIP2;5 monomers. The helices of TM1 and TM2 are shown in green and pink for ZmPIP1;2 and ZmPIP2;5 respectively. Residues occurring at the monomer-monomer interface that are different between ZmPIP1;2 and ZmPIP2;5 are labeled and are shown in space-filling representation.

**Figure 6:**
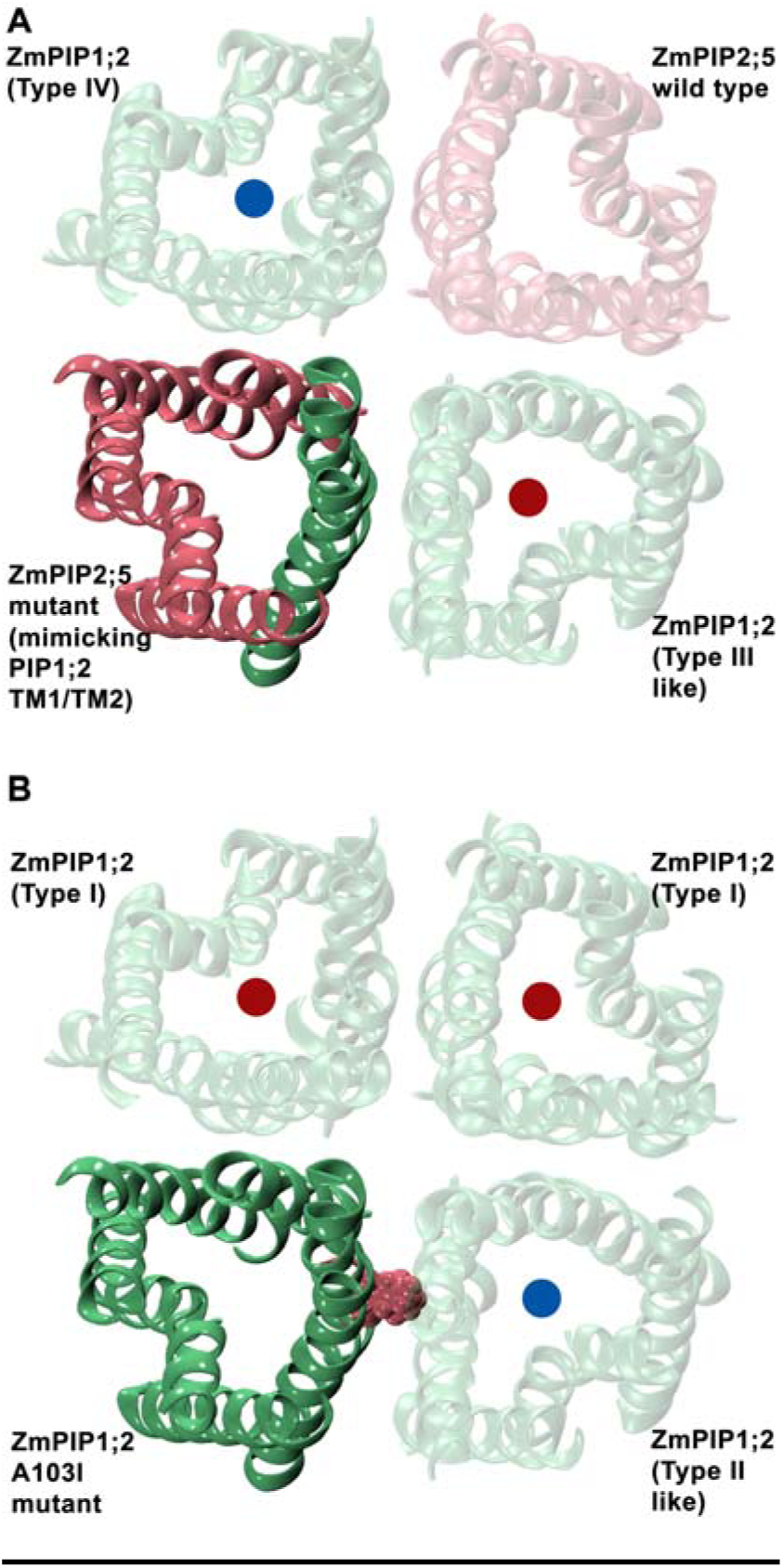
Initial configuration of two mutant systems. (A) One of the ZmPIP2;5 monomer (solid pink) in the 2:2(d) hetero-tetramer was mutated at 7 positions to mimic ZmPIP1;2 so that the adjacent ZmPIP1;2 monomer will have Type III-like interface. The mutated helix in ZmPIP2;5 is shown in solid green. (B) ZmPIP1;2 homo-tetramer with a single mutation of A103I at the TM2 position in one ZmPIP1;2 (solid green) is shown. All the wild-type monomers are shown transparent in green (ZmPIP1;2) or pink (ZmPIP2;5). The red and blue dots show the expected water permeation behavior of the corresponding monomers, with the red dot (Type I/Type III interface) indicating low expected permeation and the blue (Type II/Type IV interface) indicating high expected permeation.

We have also plotted the temporal channel radius profiles of both ZmPIP1;2 monomers in the mutant 2:2 (d) system (Figure S4 in Supporting Information). While the Type IV ZmPIP1;2 shows slight constriction in the ar/R selectivity filter region, the Type III-like ZmPIP1;2 is occluded right from the extracellular side up to the ar/R selectivity filter. This explains the poor conductivity of the Type III-like ZmPIP1;2 monomer. The MD simulations of the system in which the interfacial residues of ZmPIP2;5 were mutated to that corresponding to ZmPIP1;2 demonstrated that the TM4-TM5 interface of ZmPIP1;2 in the tetramer has a major role in modulating the water transport in ZmPIP1;2 channels.

### Contacts between residues at the monomer-monomer interface

To investigate the structural basis for increased water permeation in ZmPIP1;2 monomers from Type II/Type IV group, we calculated the contacts between heavy atoms of the interacting monomers that are within 4 Å distance and contact maps were generated as described in the Methods section. Interfaces formed by both the TM4-TM5 and TM1-TM2 regions of a monomer were considered for this purpose. Contact maps for the ZmPIP1;2 monomers in homotetramers (4:0) and ZmPIP1;2 monomers from Type II/Type IV category were generated (Figure S5 in the Supporting Information) and were used to obtain the difference maps between the contacts formed by the two ZmPIP1;2 groups mentioned above (Figure S6 in the Supporting Information). The most striking feature observed in the difference map obtained from ZmPIP1;2 homo-tetramer and the hetero-tetramer in which ZmPIP1;2s have the Type II/Type IV interface is due to the contacts formed by the residue Ile-93 from TM2 of ZmPIP2;5 (Figure 7A). This residue is substituted by Ala (the equivalent position is Ala-103) in ZmPIP1;2. Ile-93 in ZmPIP2;5 participates in multiple contacts with the TM4-TM5 region of ZmPIP1;2 in the monomer-monomer interface. It shows enhanced interactions with Ile-190, Thr-194, Phe-220 and Ala-221 from the TM4-TM5 region of ZmPIP1;2 (Figure 7A). Such extensive interactions are not possible for the small residue Ala. Based on the mutation studies, Yoo et al. ^50^ have concluded that Ile-93 is important to stabilize the TM5 conformation at the monomer-monomer interface. However, they studied the importance of residues in tetramer formation of AtPIP2;1 homotetramers. The role of I93 in influencing water transport in the hetero-tetramer environment remains to be elucidated.

**Figure 7:**
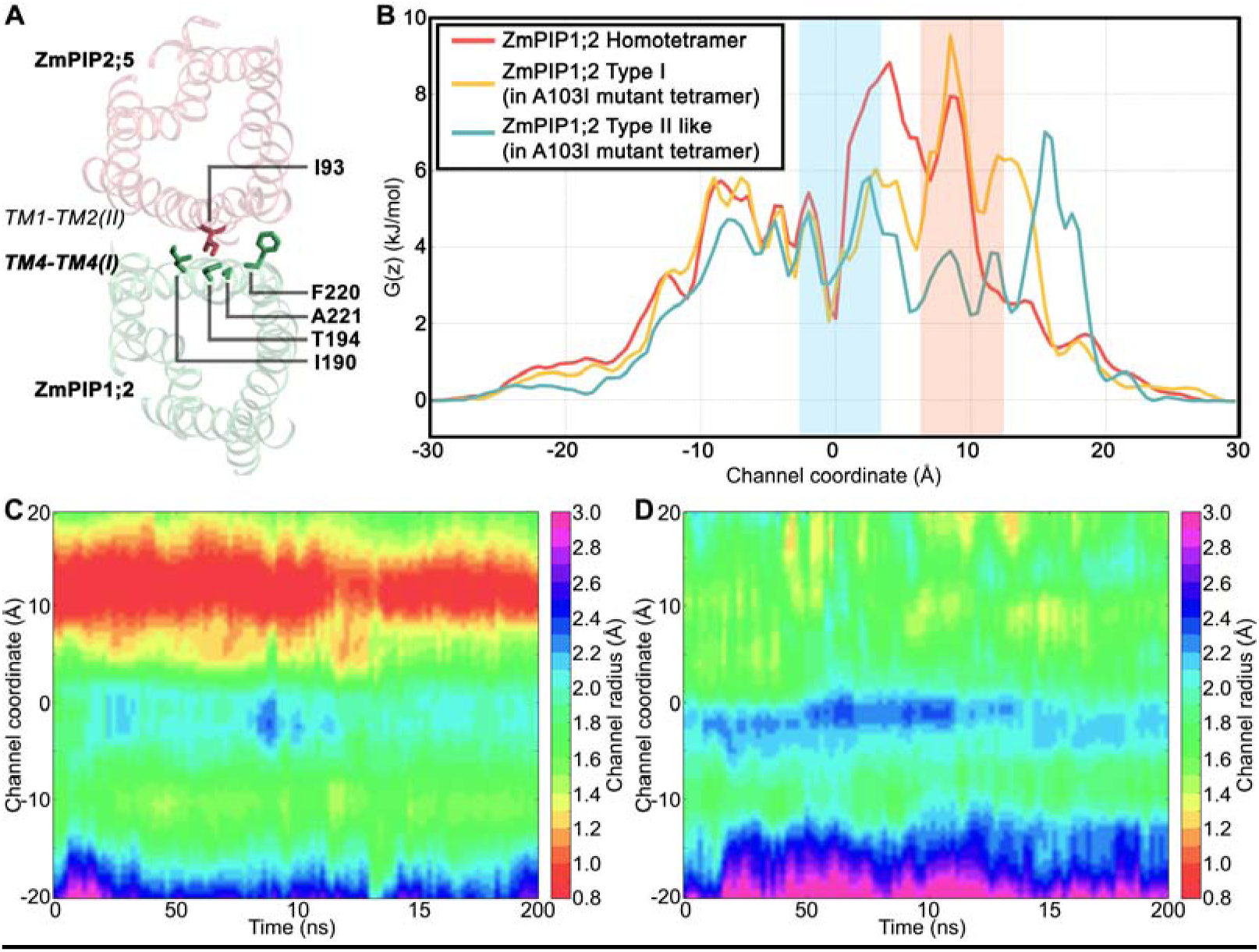
(A) The residue I93 from TM2 of ZmPIP2;5 at the monomer-monomer interface exhibit enhanced interactions with several bulky residues (I190, I194 and F220) from TM4 of ZmPIP1;2 in hetero-tetramers belonging to Type II or Type IV category. Interacting residues are labeled and are displayed in stick representation. Only two of the interacting monomers in the tetrameric system are shown. (B) Average PMF profiles calculated for monomers of ZmPIP1;2 homo-tetramers, ZmPIP1;2 monomers of Type I category from the A103I mutant homo-tetramers and ZmPIP1;2 monomer which acquired Type II-like interface in the mutant homo-tetramer. Temporal channel radius profiles of (C) Type I ZmPIP1;2 monomers and (D) ZmPIP1;2 monomer with Type II-like interface from two independent simulations of the mutant homo-tetramer systems.

To test the importance of Ile-93, we mutated Ala-103 in one of the ZmPIP1;2 monomers of the wild-type homotetramer to Ile (Figure 6B). We performed two independent simulations using the same protocol as described in the Methods section. We examined the water permeability and calculated the *p_f_* values. We compared the transport properties of ZmPIP1;2 monomer whose TM4-TM5 region interacts with the TM1-TM2 of mutant ZmPIP1;2. Our results show that this ZmPIP1;2 monomer in contact with the A103I mutant ZmPIP1;2 resulted in an average number of 226 water permeation events in the two simulations and a mean osmotic permeability of 6.2. In comparison with this monomer, the other wild-type monomers transported only 4 to 15 water molecules with the *p_f_* values ranging from 1.04 to 1.61. This is very similar to the ZmPIP1;2s in the wild-type homotetramer system which transported, on an average, only 14 water molecules. To further consolidate the conclusions from this study, we have plotted PMF profiles of ZmPIP1;2 monomers from homotetramers and the three ZmPIP1;2s from the mutant homotetramers (Figure 7B) along with their temporal channel radius profiles (Figure 7C and Figure 7D). PMF profiles reveal that the large energy barrier found in the ZmPIP1;2s of homotetramers and the two Type I category ZmPIP1;2s of the mutant homotetramers at the ar/R selectivity filter region has drastically decreased in the ZmPIP1;2 whose TM4-TM5 forms the interface with the mutant ZmPIP1;2 in the mutant tetramer. The above observations are supported by the temporal channel radius profiles of ZmPIP1;2 monomers in the A103I mutant homotetramer. The two ZmPIP1;2 monomers with Type I interface show severe constriction near the ar/R selectivity filter (Figure 7C) similar to that observed in ZmPIP1;2 wild-type homotetramer (Figure 2A). However, the ZmPIP1;2 monomer whose TM4-TM5 region is in contact with the A103I mutant ZmPIP1;2 does not exhibit any such behavior and its ar/R selectivity region is wider (Figure 7D) enabling this monomer to transport more number of water molecules similar to other Type II monomers. In summary, we have elucidated that a single mutation from Ala103 -> Ile at the monomer-monomer interface has a dramatic effect on the water permeation of the neighboring monomer with which it interacts at the monomer-monomer interface. Our in silico mutation studies have thus identified a key residue in ZmPIP2;5 which may be sufficient to enhance water permeation of neighboring ZmPIP1;2 monomers in a hetero-tetramer.

### Essential dynamics of homo- and hetero-tetramers of PIP1 and PIP2

Essential Dynamics analysis, also known as Principal Component Analysis (PCA), captures the dominant modes of protein motion by transforming the high-dimensional protein motions into low-dimensional representations. Essential dynamics analysis of homotetramers and hetero-tetramers of ZmPIPs was performed as described in the Methods section to understand the factors at the molecular level that lead to ZmPIP1;2s in hetero-tetramers conducting high levels of water molecules. Results of MD simulation studies clearly implicated the monomer-monomer interface involving TM4-TM5 of ZmPIP1;2 for the high water permeability. To identify the biologically important motions in the PIP tetramers, we performed PCA on the ZmPIP1;2 monomers from homo-tetramers and ZmPIP1;2 monomers that belong to Type II/Type IV category in the heteromeric systems. Figure S7 (See Supporting Information) shows the cumulative contribution of eigenvectors to the overall motion of the protein. Eigenvectors describe the collective motions of the atoms and a small number of them are responsible for most of the protein dynamics. About 80% of the total mean square fluctuations can be described by the first 10 eigenvectors and the first eigenvector alone represents more than 40% of the total fluctuations. We visualized the direction and the extent of fluctuation in TM5 of ZmPIP1;2 using the first eigenvector (Figure 8).

**Figure 8:**
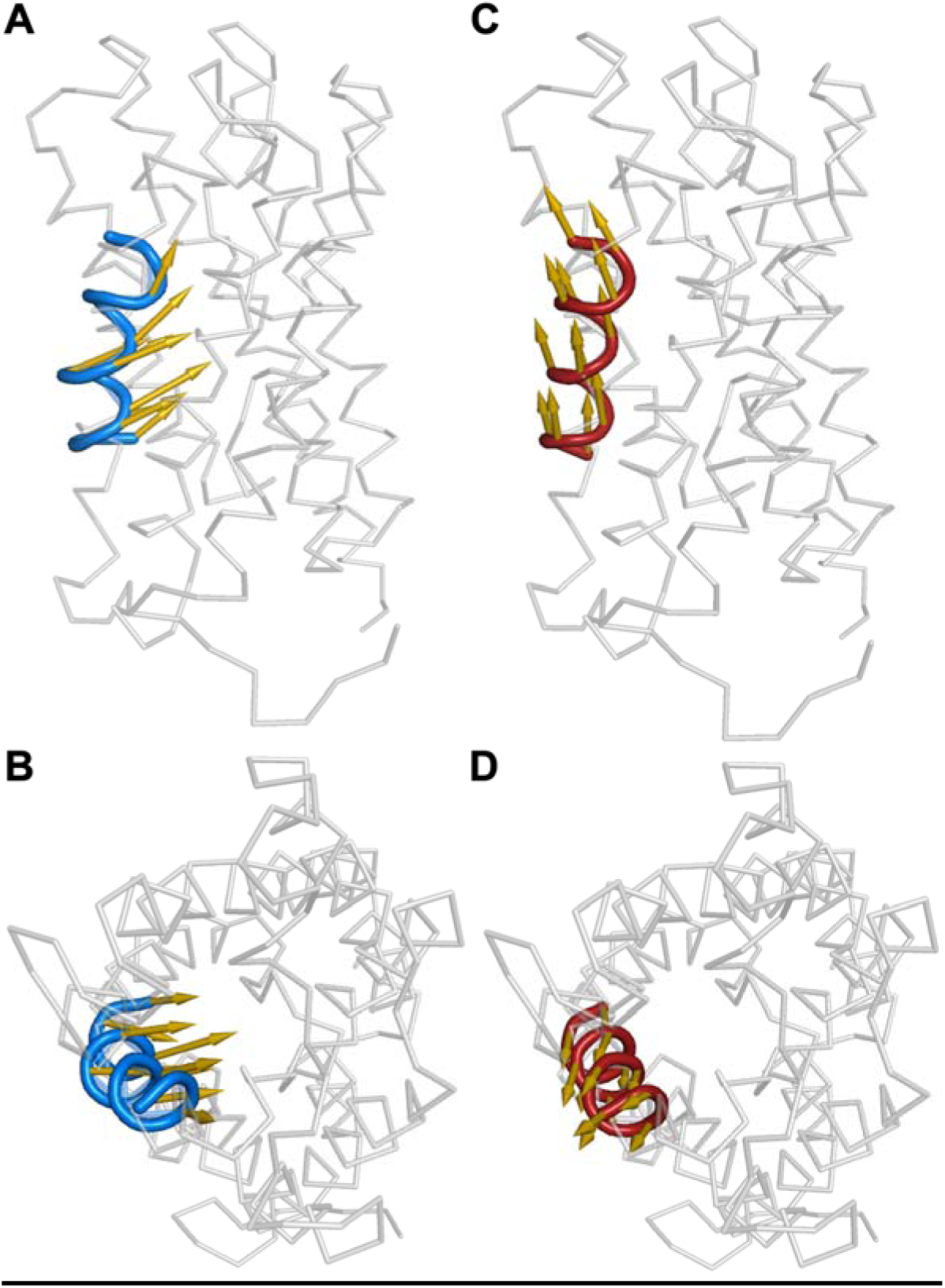
First eigenvectors, shown as arrows, depicting the direction of motion of TM5 (blue) for ZmPIP1;2 monomers from homo-tetramers viewed (A) parallel to the membrane and (B) down from the extracellular side. The direction of first principal component shown for TM5 (red) from ZmPIP1;2 monomers belonging to Type II/Type IV category viewed (C) parallel to the membrane and (D) down from the extracellular side.

In the case of ZmPIP1;2s in homo-tetramer, the movement of TM5 is towards the center of the channel (Figure 8A and 8B) while the TM5 of ZmPIP1;2 from Type II/Type IV group moves in parallel to the channel axis towards the extracellular side (Figure 8C and 8D). This could explain why the channel radius profiles of ZmPIP1;2s from homo-tetramers exhibit a pronounced constriction resulting in a narrow ar/R selectivity filter which makes the ZmPIP1;2s in homo-tetramers poor water-conducting channels. However, we do not see the same behavior in ZmPIP1;2s from Type II/Type IV group in which the TM4-TM5 regions have strong interactions with the TM1-TM2 region of ZmPIP2;5. Thus the essential dynamics analysis has helped to elucidate reasons for high water transport of ZmPIP1;2s that belong to Type II/Type IV category.

## Discussion

Oligomerization of water channels and glycerol facilitator and the role of specific residues have been investigated for several members of MIP channel family including AQP4, AQP11, GlpF and AQPcic ^76–79^. However, when plant aquaporins like maize ZmPIPs form heterotetramers, then they can interact with two different adjacent monomers. ZmPIP1;2s in hetero-tetramers have two interfaces, one involving the TM1-TM2 helical segments and the other formed by TM4-TM5 transmembrane helices. Previous experimental studies have reported the increase in water transport in PIP1s when they are part of hetero-tetramers with PIP2s ^80^. Karina Alleva and her colleagues investigated the tetramer formation of BvPIP1;1 and BvPIP2;2 from *Beta vulgaris* and found the formation of hetero-tetramers with different stoichiometries (3:1, 2:2 and 1:3) localized in the plasma membrane ^49^. Further functional studies demonstrate that all hetero-tetrameric configurations coexist and exhibit similar efficiency in transporting water molecules. They concluded that the formation of homo- and hetero-tetramers can modulate the water permeability of the cell plasma membrane. Yoo et al. ^50^ carried out extensive mutagenesis studies on *Arabidopsis* PIP2;1 to identify residues that could play important role in tetramer formation. Their studies indicated that residues in TM1, TM2 and TM5 involved in transmembrane helix-helix interactions within and between monomers are critical in tetramer assembly. Chaumont and his colleagues used a model building approach to identify the residues in hetero-tetramer that are involved in interactions with adjacent monomers and these residues were subjected to mutation studies ^51^. Residues that inactivate the PIP1/PIP2 channel within homo- or hetero-tetramer were identified. However, the functional and mutational studies did not investigate the possible role of two different interfaces of PIP1 in hetero-tetramer environment. When present in a hetero-tetramer, PIP2 can form an interface from either of the two PIP1 sides. It was not envisaged that the two interfaces will behave differently and experimental studies did not distinguish between the two interfaces. Our extensive MD simulation studies on homo- and hetero-tetramers with different combinations of ZmPIP1;2 and ZmPIP2;5 monomers clearly elucidated that the interface with TM4-TM5 segment of ZmPIP1;2 has the greatest influence on water permeation properties. When TM4-TM5 of ZmPIP1;2 interacts with the TM1-TM2 of ZmPIP2;5, the amount of water transported through the ZmPIP1;2 channel is at least three to four-fold higher than that found for the ZmPIP1;2s when they are present in the homo-tetramers. This has been observed repeatedly in our studies in ZmPIP1;2s that are from Type II/Type IV category with different stoichiometries. However with the TM1-TM2 interface of ZmPIP1;2s, even when it is interacting with the TM4-TM5 helices of ZmPIP2;2, no such increase in water transport is observed. This was clearly seen in ZmPIP1;2 members from Type I or Type III group in our simulations. To further elucidate the factors responsible for high water transport in ZmPIP1;2s from Type II/Type IV group, we generated *in silico* mutants replacing the ZmPIP2;5 residues at the interface to that found in ZmPIP1;2 monomers. In this experiment, seven residues of ZmPIP2;5 at the PIP1-PIP2 interface were substituted by that found in ZmPIP1;2. The number of water permeation events showed a dramatic decrease in the mutant tetramer. Using contact map analysis, we then identified a single residue I93 in ZmPIP2;5 that makes the maximum number of contacts with the adjacent ZmPIP1;2 monomer. The equivalent residue for I93 in ZmPIP1;2 is Ala. When Ala in ZmPIP1;2 was mutated to the corresponding residue in ZmPIP2;5 which is Ile, the TM4-TM5 of neighboring ZmPIP1;2 interacted with increased number of contacts with the mutant ZmPIP1;2. As a result, this mutation resulted in higher water transport of ZmPIP1;2 clearly indicating a gain-of-function role for this specific mutation. We conclude that the TM4-TM5 interface of ZmPIP1;2 in hetero-tetramer environment can greatly modulate the water transporting properties by interacting either with TM1-TM2 of ZmPIP1;2/ZmPIP2;5. Depending upon its interactions with PIP1/PIP2, the amount of water that can be transported can be regulated.

## Conclusions

The biological significance of tetramer formation in the superfamily of major intrinsic proteins is an intriguing phenomenon that has been investigated experimentally. Although each monomer in the tetramer is a functional channel, the important question is whether the adjacent monomers have any influence on the transport properties of an individual channel within the tetramer assembly. Plant plasma intrinsic proteins (PIPs) have been investigated using functional and mutational studies to address this specific issue. Although the PIP1s and PIP2s show very high sequence identity, the transport properties of PIP1s differ significantly from that of PIP2s. While PIP1 homo-tetramers are considered functionally inactive in some plant species, PIP2s are shown to be efficient water channels. However when present as part of hetero-tetramers, PIP1s become active and are involved in abundant water transport. What makes this transition for a PIP1 channel from inactive in a homo-tetramer to efficient water channels in hetero-tetramers has been the focus of many researchers. In the current study, we have carried out molecular dynamics simulations of homo- and hetero-tetramers of ZmPIP1;2 and ZmPIP2;5 with different stoichiometries and configurations and analyzed different properties including water permeation events, temporal channel radius profiles and PMF profiles. We focused on the two interfaces of ZmPIP1;2 formed by TM4-TM5 and TM1-TM2 helical segments. Their influences on water transport in both homo- and hetero-oligomers were examined. We clearly demonstrate that the interface formed by TM4-TM5 with ZmPIP2;5 in hetero-tetramer shows the greatest effect and results in dramatic increase in the number of water permeable events. The interface of ZmPIP1;2 formed by TM1-TM2 with ZmPIP2;5 does not produce the same effect and behaves very similar to PIP1s in homo-tetramers. Our MD simulation studies have offered a molecular explanation for the high water conductivity of PIP1s in hetero-oligomeric environment. We further demonstrated that the interface residues of ZmPIP2;5 in hetero-tetramer when substituted by those found in the equivalent positions in ZmPIP1;2 again resulted in reduced water transport. Furthermore, we have specifically identified a single residue, Ala-103, in ZmPIP1;2 which when mutated to the corresponding ZmPIP2;5 residue, Ile, has markedly increased water transport in the adjacent ZmPIP1;2 channel. In other words, the interaction of ZmPIP1;2 residues at the TM4-TM5 interface with ZmPIP2;5 residues, the ZmPIP1;2 channels become active water transport channels. Essential dynamics analyses indicated important differences between hetero- and homo-tetramers in the movement of TM5 of ZmPIP1;2. We have thus demonstrated that the two monomer-monomer interfaces of ZmPIP1;2 are not the same and the TM4-TM5 interface when interacting with ZmPIP2;5 in hetero-oligomers is responsible for higher water transport and thus can modulate the water transporting properties of ZmPIP1;2s. Further experimental studies combined with simulation studies can help to understand whether this phenomenon is applicable to all PIPs from within the same species. Additionally, the universality of the role of TM4-TM5 in modulating the water transporting properties of PIP1s for all the plant species remains to be established.

## Acknowledgements

We gratefully acknowledge the High Performance Computing Facility at IIT-Kanpur. We thank all our lab members for useful discussions.

